# The critical role of spatio-temporal control in combinatorial chemo- and magnetic hyperthermia thermo-therapy: ‘the where’, ‘the how’ and ‘the when’

**DOI:** 10.1101/2023.06.15.545102

**Authors:** Lorena García-Hevia, Andreia Patrícia Magalhães, Nuria Genicio, Íñigo Casafont, Milene Costa da Silva, Mónica López Fanarraga, Manuel Bañobre-López, Juan Gallo

## Abstract

Combinatorial treatments hold the key to the future of cancer treatment as they enhance therapeutic indexes by inducing synergistic effects and reducing resistance processes, while often providing a safer option for patients with fewer off-target effects. However, combinatorial treatments bring extra problems to cancer management not only derived from the actual compatibility of the treatments, but also from their hands-on administration. Operational parameters such as administration order and dosing (dose, spacing) have to be optimized in order to positively impact patient prognosis. Here we present a systematic study on the optimization and the importance of these parameters within the framework of a combinatorial chemo-/thermo-therapy. Parameters like ‘where’, ‘how’ and ‘when’ are investigated in detail. Furthermore, we delve into the underlying biochemical mechanisms driving the observed effects through transcriptome analysis.

---

Cancer continues to be one of the main causes of death in developed countries and, even though recent therapeutic advances have brought hope to many patients, these developments are punctual rather than having a wide application. For instance, while the CAR-T cell technology has demonstrated exceptional efficacy in certain types of blood malignancies, their performance is relatively modest when it comes to treating solid tumors.^1, 2^ One of the strategies that is showing more promise is the combination of several therapeutic modalities to treat the condition. The idea of attacking a tumor from several fronts appears due to the propensity of tumor cells, and particularly tumor stem cells, to develop resistance mechanisms against treatments. This problem is more obvious in chemotherapy, where many different resistance mechanisms have been described (drug inactivation, target alteration, enhanced DNA repair, cell death inhibition, decreased drug uptake, drug efflux…),^3, 4^ but it is present in most other forms of therapy (e.g. radio-, immuno-therapy).^5, 6^ The possibilities for treatment combinations are vast, ranging from the integration of various chemotherapeutic drugs targeting different molecular pathways, to the synergistic combination of entirely distinct cancer treatment modalities (either well established - chemo-/radio-/immuno-therapy - or more novel ones - PDT, thermo-therapy). However, the administration of combinatorial treatments brings extra concerns associated with the compatibility of the operational parameters of each treatment option (administration order, doses, technical parameters…) and the practical design of the actual administration protocol.

In addition to resistance mechanisms, specificity plays a crucial role in determining the effectiveness of cancer treatments. Targeted drugs have made a profound impact on patient prognosis and breast cancer serves as a notable example. Over the years, the prognosis for molecular subtypes such as Luminal A (ER+, PR+, HER2-), Luminal B (ER+, PR+, HER2+/-), and HER2+ (ER-, PR-, HER2+) has significantly improved. However, the Triple-negative subtype (ER-, PR-, HER2-) continues to present significant challenges in terms of survival rates.^7, 8^ These differences come, undoubtedly, from the introduction in the clinic of targeted therapies tailored for the Luminal and HER2+ subtypes (in the form of endocrine therapy and drugs like trastuzumab or lapatinib), whereas no specific treatment is still available for the Triple-negative subtype.

The introduction of nanotechnology and nanomedicine into the oncology arena is meant to increase the therapeutic options available for cancer patients.^9, 10^ Nanostructured drug delivery systems (DDSs) are ideal systems to combine different therapeutic drugs or even modalities into a single entity, easing administration and dosing. The different pharmacodynamic (PD) and pharmacokinetic (PK) properties of drugs, for example, are partially disregarded when co-encapsulated into a single nanosystem as then only one entity is delivered, thereby neglecting the individual properties of each drug. On top of this, hybrid organic-inorganic nanocomposites enable the combination of drug delivery (single drug or co-delivery) through the organic counterpart, with other treatment modalities mediated or enhanced through the inorganic component.^11–13^ Hybrid nanocomposites incorporating magnetic nanostructures^14, 15^ are promising candidates for combinatorial treatment administration. In addition to the drug delivery capabilities mentioned earlier, these hybrid magnetic nanocomposites offer the advantages of thermo-therapy and externally triggered drug release through magnetic hyperthermia (MH). Moreover, in a properly designed system, the intrinsic magnetic properties of the hybrid nanocomposites enable the use of magnetic resonance imaging (MRI) for non-invasive, quasi-real-time monitoring of treatment progress. While nanocomposite systems can streamline pharmacokinetics/pharmacodynamics (PK/PD) by consolidating multiple behaviors into a single framework, other administration parameters must be meticulously optimized to maximize therapeutic indexes. Therefore, precise control of the application scheme (including timing, dose, location and operational parameters) is crucial for ensuring the efficacy of the combination therapy.

In this report we present an in-depth study of a lipid-based nanocomposite (magnetic lipid nanovehicles, mLNVs) co-loaded with a clinical chemotherapeutic agent (doxorubicin, DOX) and magnetic nanoparticles (MNPs), as a combinatorial chemo-/thermo-therapeutic agent against melanoma. In this system, MH rather than being used primarily to directly thermally ablate cancer cells, has its main role as an external trigger to induce a change in the release profile of DOX. In this role, MH enhances specificity being as it is a local treatment driven by the controlled spatio-temporal application of an external alternating magnetic field. The operational parameters and interrelationships between both forms of treatment are explored in detail to optimize the therapeutic outcomes. Finally, the biochemical processes behind the superior performance of the combined treatment versus its individual counterparts, are explored through transcriptome analysis.

## RESULTS and DISCUSSION

The preparation and characterization of the mLNVs and mLNVs-DOX were described in detail before. ^16^ mLNVs are lipid-based magnetic nanocomposites that use a natural wax (carnauba wax) as a matrix to co-encapsulate DOX, as chemotherapy, and 10 nm magnetite (Fe3O4) nanoparticles (Table S1). Tween 80 is used as a surfactant to control the size and distribution (204 ± 4 nm) of these mLNVs, as well as to stabilize them in aqueous solution. All the components of these nanocomposites are FDA approved for human use. Beyond physicochemical characterization, the functional properties of these hybrid magnetic nanocomposites are very promising for biomedical applications such as MRI (transversal relaxivity = 249 mM^-1^ s^-1^ at 37 °C and 3.0 T) and MH (SAR = 517 W g^-1^ at 869 kHz and 25 mT) enabling their theranostic application. In oncology, traditional MH strategies involve raising the temperature of tumor tissues to or above 42-45 °C, exploiting the limited thermo-regulation ability of cancer cells.^17^ This strategy is limited by the high concentration of magnetic probes required locally to reach the desired temperature. So far, such high concentrations can only be reached through direct intratumoral injection of highly concentrated magnetic nanoparticle dispersions, which has limited its translation into the clinic to the treatment of brain tumors.^18^ In our design, direct tumor thermal ablation is only contemplated as a collateral (but positive) effect, while MH is primarily used to control the release profile of the drug from the mLNVs.^19^ This means that in this chemo/thermo combinatorial treatment, both thermo and chemo are very closely linked, and their operational parameters have to be carefully optimized. Additionally, the clinical application of DOX, although first-line treatment for different types of cancer, is limited by the activation of resistance-related mechanisms^20, 21^ and the potential for cardiotoxicity,^[23, 24]^ which can result in poor prognosis and survival.^21–27^ Compared to conventional “free” drugs, nanoparticle-based DDSs have the potential to improve the stability, solubility and biocompatibility of the encapsulated drugs, showing higher bioavailability, enhanced permeation and retention effect and precise targeting.^28, 29^ Nevertheless, it is essential to optimize the local activity of DDSs by enabling on-demand drug release and achieving effective accumulation at the target site. This optimization plays a vital role in the success of the therapy and contributes to improving the post-treatment quality of life for patients by minimizing unwanted toxicities and enabling a broader range of dosages.

### ‘The Where’

MH is a local treatment, meaning that mLNVs located within the alternating magnetic field (AMF) will generate a temperature increase proportional to their concentration, as well as the frequency and the magnitude of this AMF. Conversely, mLNVs farther away from the AMF will not induce any temperature change. This local character of MH can help overcome the lack of molecular specificity of the mLNVs. To illustrate this effect, a bespoke sample holder (Fig S1A and B) was fabricated to be used in combination with an AMF applicator designed to work specifically with *in vitro* models, maintaining both temperature (37 °C) and atmospheric (5 % CO_2_ and high humidity) conditions (Fig S1C). This system uses a planar coil (68 mm external diameter, 8 turns) placed right on top of the sample,^30^ and allowed us to evaluate the effect of the combined chemo-/thermo-therapy on cell viability as a function of the distance from the center of the AMF coil. Melanoma cells were seeded in 25 mm glass covers, placed on the holder and submitted to 1 h of AMF right after the administration of the mLNVs-DOX. After live cell staining with calcein, a series of confocal images (Fig 1A) showed that in the coil center (left of the image) the density of viable cells was significantly reduced compared to the control without MH (Fig 1B). As we move further away from the center of the coil, the viability gradually recovers, and at around 4 cm from the center of the coil the cell density is comparable to the control without MH. This distance, 4 cm, matches well the radius of the coil, 3.4 cm, and illustrates well the local character of the MH process.

**Figure 1.**
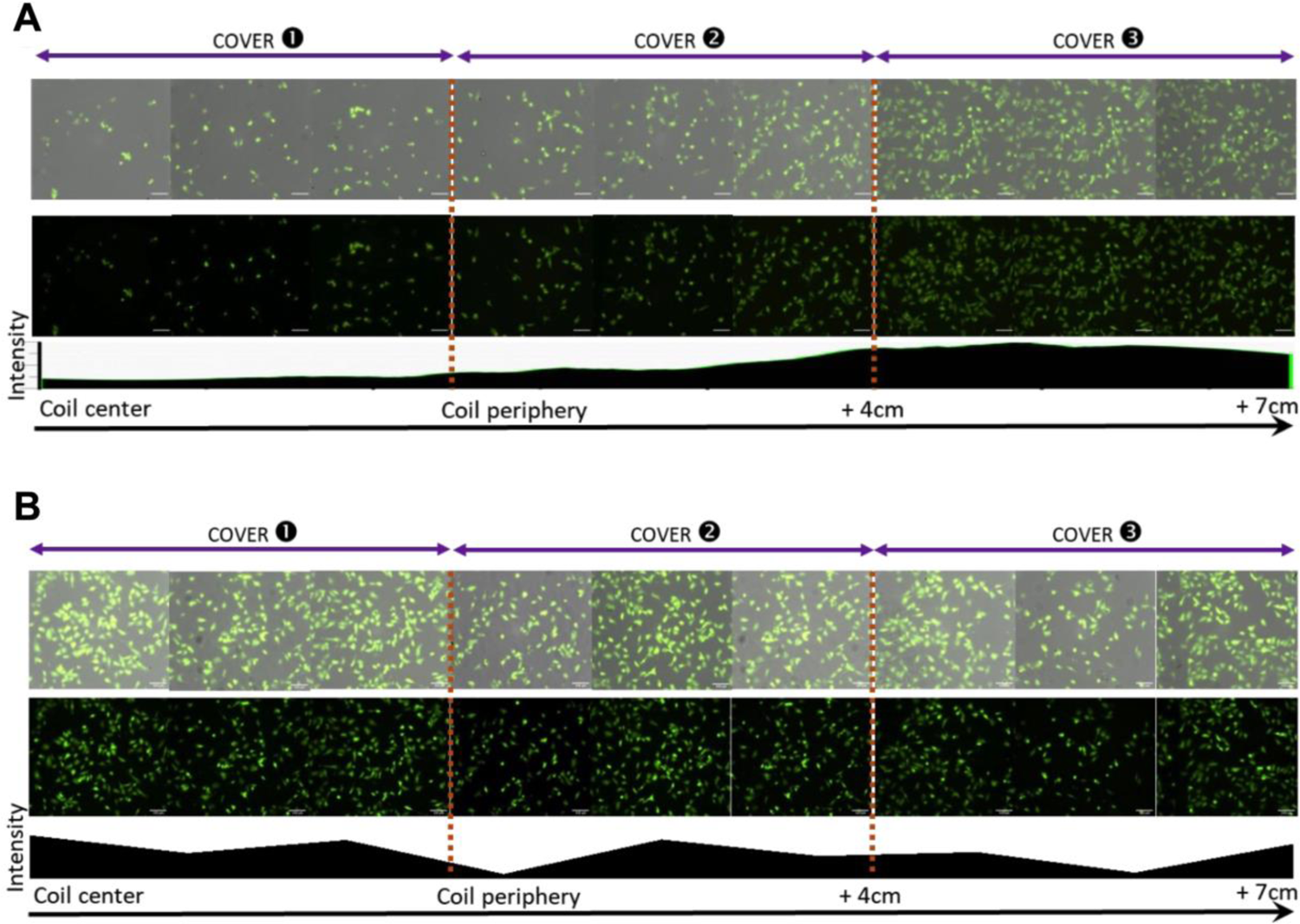
Proximity to the coil center reduces cell viability upon MH. Composition of confocal images of live-stained (calcein staining) melanoma cells incubated with mLNVs-DOX (0.5 μg DOX/mL) and subjected (**A**) or not (**B**) to MH for 1h. Bottom row, quantification of the fluorescence intensity along the different coverslips; Middle row, confocal images (green channel) of the different coverslips; Top row, brightfield plus confocal images (green channel) of the different coverslips.

### ‘The How’

Once the magnetic effector (in this case the mLNVs) is selected, the effect of MH depends on the selected frequency, field intensity and application time of the AMF. Setting time aside (in a final clinical application this parameter will be defined not only by the efficiency of the whole process but also by patient comfort), MH effect depends more precisely on the frequency and the square magnetic field intensity of the AMF. Consequently, it is more effective to adjust the field intensity rather than the frequency, as the anticipated differences in heat generation would be more significant. Therefore, in order to assess the impact of MH operational parameters on the combined chemo-/thermo-therapy efficacy, the AMF intensity was then varied as shown in Fig 2A. As predicted, changing the specifications of the magnetic field had a clear impact on the final viability of melanoma cells after the chemo-/thermo-therapy. Under the conditions used before (20 mT at a mLNVs-DOX concentration of 0.5 μg DOX/mL), melanoma cell viability was reduced to around 26 %. By reducing the magnetic field to 10 mT, the viability increased to values over 50 %. An additional relevant operational parameter that deserves attention is the application mode. While continuous application is typically preferred, a few reports mention pulsed application schemes with potentially superior performance over continuous schemes.^31^ To test the effect of the application mode on the cell viability, our standard protocol (224 kHz and 20 mT at a mLNVs-DOX concentration of 0.5 μg DOX/mL for 1 h) was compared with a pulsed protocol (224 kHz and 20 mT at a mLNVs-DOX concentration of 0.5 μg DOX/mL for 3 cycles of 10 min with 20 min of rest in between pulses). The tested pulsed protocol, as shown in Figure 2B and 2C, did not demonstrate any improvement compared to the continuous protocols. In fact, the resulting cell viability under these conditions (64%) was similar to the viability achieved when reducing the applied field to 10 mT (55%).

**Figure 2.**
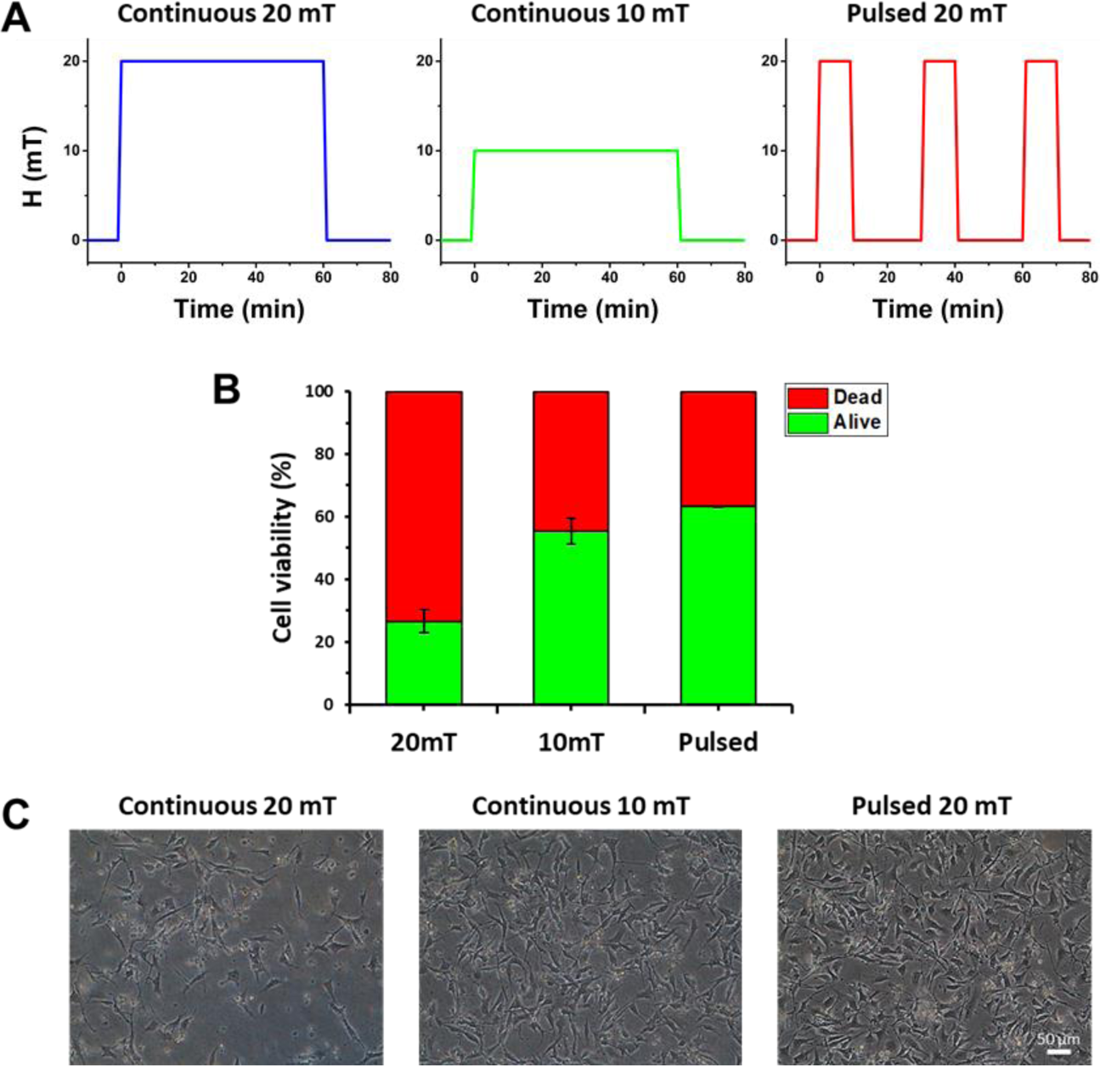
**(A**) Different magnetic field (H) conditions used to test ‘the How’. (**B**) Cell viability of melanoma cells after administration of mLNVs-DOX (0.5 μg DOX/mL) and application of AMF either at 20 mT for 1h, 10mT for 1h or 20 mT for 10 min x3 with 20 min rest in between. (**C**) Bright field microscope images of the cell cultures after the protocols described in B.

### ‘The When’

Probably more interesting than the foreseeable local effect of chemotherapy-combined MH and the predictable influence of AMF operational parameters, is the interrelationship between mLNVs-DOX pre-incubation time before MH application and the final viability outcome. As previously stated, MH offers an additional level of control over the release of DOX from mLNVs. Consequently, the temporal control of the process emerges as a crucial factor in the overall therapy. A premature application of MH can result in the extracellular release of the drug, leading to final cell viability comparable to the one provided by the free drug. On the other end, a late MH application would not affect drug release (most of the drug would have already been released) and the final effect on the viability would be similar to the one obtained in the absence of MH. Therefore, to investigate the role of time in this context, an experiment was designed following the scheme depicted in Fig 3A.

**Figure 3.**
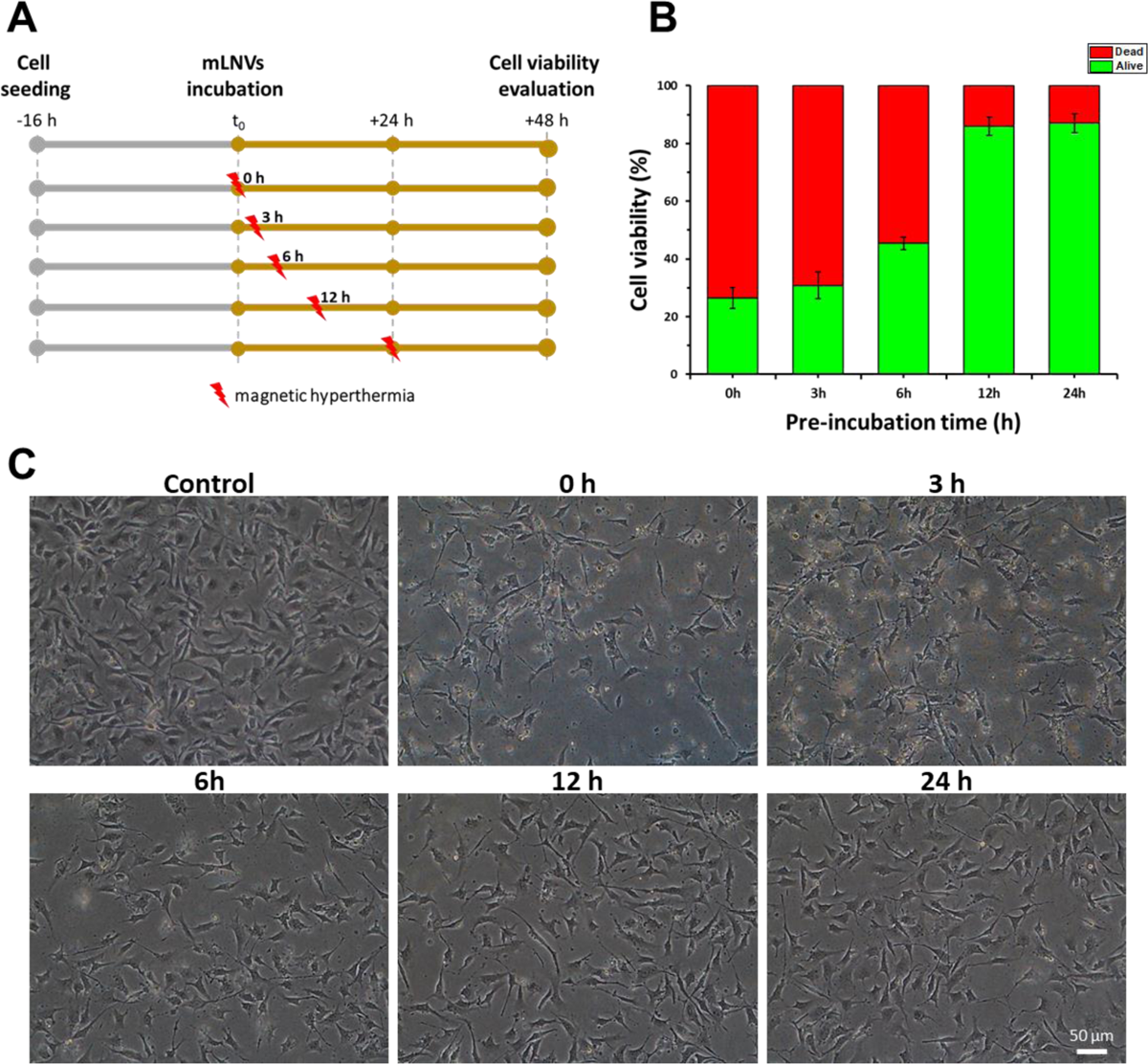
(**A**) Experimental design for the *in vitro* chemo-/thermo-therapy pre-incubation time effect evaluation. (**B**) Melanoma cell viability after pre-incubation with mLNVs-DOX (0.5 μg DOX/mL) for 0, 3, 6, 12 and 24 h and continuous application of AMF at 20 mT for 1h. (**C**) Phase contrast microscope images of the cell cultures after the protocols described in A.

The effectiveness of chemo-/thermo-therapy was found to be significantly superior at short pre-incubation times (up to 6 hours) compared to long pre-incubation times (12-24 h) (Fig 3B). Indeed, cell death inversely correlates with pre-incubation times even at the shortest time tested (0 h pre-incubation plus 1 h AMF application), when the effect is maximum, reaching values below 30% of living cells. When mLNVs-DOX were pre-incubated with the melanoma cultures for 3 h, the effect on cell viability was still strong and very similar (∼30%). At 6 h pre-incubation, cell viability increased to ∼50 % and at 12 and 24 h pre-incubation times the effect of the chemo-/thermo-therapy is negligible (viability around 90 %). The results from these longer pre-incubation time points are in agreement with cell viability results when mLNVs-DOX are applied without MH (negligible effect, Fig S2). To further corroborate this effect, the release profile of DOX from the mLNVs-DOX under passive conditions at 37 °C was characterized (Fig S3A and B). The results showed that the release rate of DOX is highest at the initial time point (0 h) and decreases exponentially over time (Fig S3C). By 12 h the speed has already slowed down and even more by 24 h, meaning that most of the DOX is already free and the release cannot be influenced by the MH process anymore. Moreover, the release of DOX is significantly faster under MH than under passive conditions, with over an 80% increase in released DOX after only 1h (Fig S3B). This, together with the fact that the encapsulation of DOX in mLNVs-DOX accelerates intracellular nuclear DOX accumulation,^32^ could explain why mLNVs-DOX together with MH reduce melanoma cell viability more efficiently at shorter pre-incubation times. This relationship holds significance in determining the final viability outcome, underscoring the importance of optimizing the timing and coordination between mLNVs-DOX pre-incubation and MH administration.

To test whether the observed enhanced effect at short pre-incubation times was dependent on operational MH parameters, the different conditions tested above to optimize ‘*the how*’ were used again with different pre-incubation times. As seen in Fig S4, a similar time-dependent effect of the pre-incubation time on cell viability is still observed regardless of the MH conditions. According to that, and to investigate whether this effect was influenced by the concentration of the chemotherapy drug, two different concentrations were also evaluated under the same MH conditions (Fig S5). The inverse correlation between pre-incubation time and cell death was clearly observed at both concentrations, of course in a clearer way with the higher concentration (1 μg DOX/mL). Interestingly, cell viability was lower at earlier time points (0, 3 and 6 h) when melanoma cells were incubated at a lower concentration (0.5 μg DOX/mL), highlighting the beneficial effects of combining thermotherapy (MH) with chemotherapy (mLNVs-DOX) for the treatment of melanoma cells.

Some reports have described the impact of MH beyond its ability to convert hysteresis losses into heat. It has been suggested that MH may exert mechanical effects on the cell membrane, which could contribute to its cytotoxic effects.^19, 33^ Although the MH conditions examined in these studies differ from the ones investigated here especially in terms of the frequencies used, for our viability tests we employed the Trypan blue exclusion assay which is a cytotoxicity test that is particularly sensitive mechanical damage as it relies on the integrity of the cytoplasmic membrane. To further investigate this possibility and to characterize the uptake of mLNVs and their intracellular distribution, melanoma cells cultures were incubated with mLNVs for different times, fixed and processed for ultra-sectioning for TEM observation. Interestingly, we found that during short incubation times of 3 h, mLNVs were predominantly observed at the cell membrane (Fig S6). However, with longer incubation times, these nanocomposites were mainly found inside the cells. This variation in the location of mLNVs over time suggests a dynamic interaction between the cells and the nanocomposites, potentially influencing their cytotoxic effects. However, further investigation is needed to gain a more comprehensive understanding of the underlying mechanisms and implications of this observation.

### Gene expression signatures

Various studies have demonstrated that hyperthermia-induced heat stress in cancer cells leads to notable alterations in gene expression. Thus, these potential changes in gene expression were investigated through transcriptome analysis to understand the underlying mechanisms and reasons for the observed synergistic effects in a combined chemo-/thermo-therapy. Additionally, as the effectiveness of the treatment can be affected by the activation of several resistance mechanisms, such as pro-survival signaling, DNA damage repair or drug efflux,^34–36^ the transcriptomic alterations related to the DOX gene resistance network were particularly examined.

Four groups were analyzed, control untreated melanoma cells, cells treated with free DOX, cells treated with mLNVs plus MH and cells treated with mLNVs-DOX plus MH. The mRNA expression analysis was performed at a 40M reads and the quality of the analysis was high (Fig S7). Genome-wide expression changes were visualized as a heatmap to identify specific genes with high fold changes and statistical significance (Fig 4A). Hierarchical clustering of the samples revealed both major divergences between the conditions and high within-group reproducibility. In addition, the heatmap analysis indicated that the distribution of the expression values was closest between control and mLNVs + MH on one side, and between free DOX and mLNVs-DOX + MH groups on the other. Overall, when cells were exposed to free-DOX, substantial changes on the expression profile of 3232 genes (1479 overexpressed; 1755 downregulated) were revealed. Almost 2000 genes showed altered expression levels in cells treated with mLNVs-DOX + MH, with approximately half of them showing upregulation and the other half showing downregulation. On the other hand, in melanoma cells exposed to control mLNVs without DOX, MH therapy only resulted in alterations in the expression of 2 genes (Fig 4B), which could be anticipated from the minor change in cell viability recorded in the cytotoxicity tests (viability ∼90%). These two genes were the *podoplanin (pdpn)* gene (overexpressed 2-fold), and the *rad9a* gene, that was downregulated (2-fold). The first gene codifies for a transmembrane receptor glycoprotein^37^ that is involved in adhesion, viability and cell cycle arrest at G2/M^38^ preventing mitosis.^39^ Interestingly, the other gene, *rad9a,* produces another cell cycle checkpoint protein, this one essential for DNA damage repair and cell cycle arrest.^40, 41^ It is interesting to note that the downregulation of *rad9a* impeding DNA repair and promoting cell division, may work in conjunction with the overexpression of *pdpn* which is heavily involved in several tumoral processes, being currently considered a biomarker associated to poor prognosis.^42^ As such, a preliminary analysis of the data suggests that the application of mild hyperthermia through mLNVs and MH might not be adviced as a ‘solo’ melanoma treatment as it can potentially contribute to an increase of the aggressiveness of the tumors. Undoubtedly, more work is be required to clarify the effect of the differential expression of these two genes in the final outcome of the therapy.

**Figure 4.**
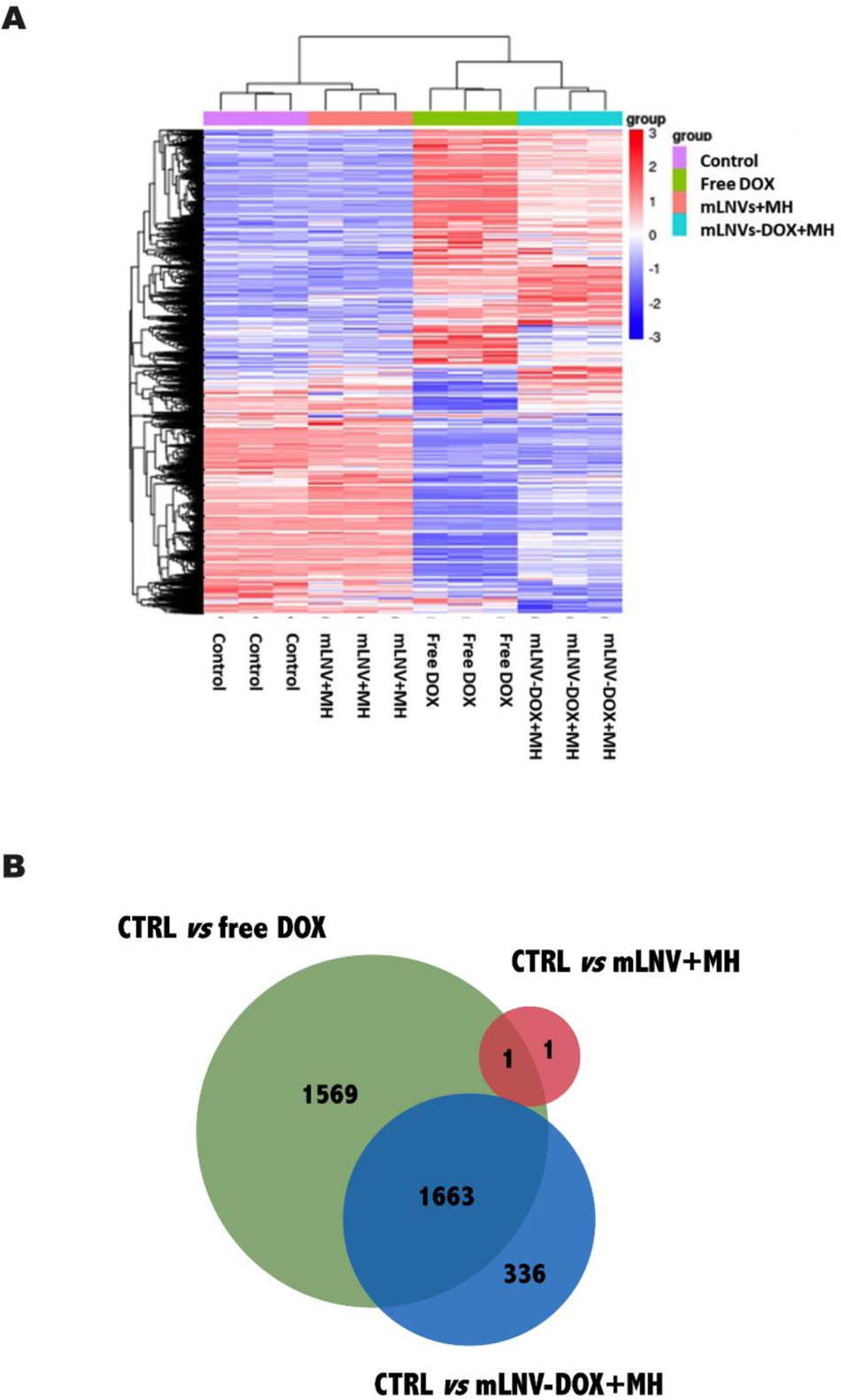
**(A)** Hierarchical Clustering Heatmap. The overall results of FPKM cluster analysis, clustered using the log2(FPKM+1) value. Red color indicates genes with high expression levels and blue color indicates genes with low expression levels. The color scale from red to blue represents log2(FPKM+1) values raging from large to small. **(B)** Venn diagram of the statistically (FC ± 2 and *p*-value < 0.05) differentially expressed genes between all the conditions compared to control. Circle size in B is only intended as a guide to the eye.

Furthermore, global comparison of differentially expressed genes (DEGs) shown in Figure 4B showed that 1663 genes were specifically expressed in the control vs free DOX and control vs mLNVS-DOX plus MH groups, and only 1 gene was shared by the control vs free DOX and control vs mLNVs-MH groups (*Rad9a*). Overall, chemotherapy has a significantly higher impact on the transcriptomic profile of melanoma cells than thermotherapy.

Table 1 summarizes the top 20 DEGs in each experimental group compared to the control cells. The Venn diagram for these data is presented in Figure S8. Of the top 20 DEGs, 11 genes were specifically overexpressed in the control *vs* free DOX and the control *vs* mLNVs-DOX plus MH group, while only 3 genes were commonly downregulated in both conditions. The overlapped up-regulated genes are associated with proliferation, tumorigenesis, and metastasis (*aifm3*^43^, *apoa1*^44^, *grp113*^45, 46^, *fam229a*^47^, *grb7*^48^) or autoimmune response (*gpr83*^49^) in different types of cancer cells. *Lhx3* was identified as an early-stage and radiosensitivity prognostic biomarker, a potential oncogene in lung adenocarcinoma,^50^ and is required to maintain cancer cell development of high-grade oligodendroglioma.^51^ Increased expression of coagulation factor XIII B (*f13b*) indicated excellent survival in glioblastoma patients in the Cancer Genome Atlas (TCGA) dataset. *Gm27196* and *1700072G22Rik* are antisense and long intergenic non-coding RNA (1700072G22Rik), respectively. The common down-regulated list comprises a pseudogene (*Gm17195*), a long intergenic non-coding RNA (*Gm26786*) and a micro-RNA (*miR-186*). *miR-186* has been shown to exhibit an inhibitory effect on cell proliferation, migration and invasion in melanoma as well as in other tumor cells.^52, 53^ Thus, this micro-RNA may serve as a potential therapeutic agent for melanoma tumors. Interestingly, none of these genes were significantly altered in cells treated with mLNVs plus MH without DOX, further supporting the need for of chemo/thermo combination. Overall, an analysis of GO terms associated with the top 20 up- or down-regulated genes revealed no enriched pathways in either overlapped or specific gene set.

**Table 1.** Experimental conditions for the MH tests.

| Experiment | Pre-incubation |  | Magnetic Hyperthermia conditions |  |  |
| --- | --- | --- | --- | --- | --- |
|  | time (h) | H (mT) | v (kHz) | t (min) | Mode |
| Pre-Incubation time | 0 | 20 | 224.0 | 60 | Continuous |
|  | 3 | 20 | 224.0 | 60 | Continuous |
|  | 6 | 20 | 224.0 | 60 | Continuous |
|  | 12 | 20 | 224.0 | 60 | Continuous |
|  | 24 | 20 | 224.0 | 60 | Continuous |
| Magnetic field | 0 | 10 | 224.0 | 60 | Continuous |
|  | 3 | 10 | 224.0 | 60 | Continuous |
|  | 6 | 10 | 224.0 | 60 | Continuous |
|  | 12 | 10 | 224.0 | 60 | Continuous |
|  | 24 | 10 | 224.0 | 60 | Continuous |
| Mode | 0 | 20 | 224.0 | 3 x 10* | Pulsed |
|  | 3 | 20 | 224.0 | 3 x 10* | Pulsed |
|  | 6 | 20 | 224.0 | 3 x 10* | Pulsed |
|  | 12 | 20 | 224.0 | 3 x 10* | Pulsed |
|  | 24 | 20 | 224.0 | 3 x 10* | Pulsed |
| Spatial control | 0 | 20 | 224.0 | 60 | Continuous |
| *In vivo* | 3 | 23 | 174.5 | 60 | Continuous |
|  | 24 | 23 | 174.5 | 60 | Continuous |

\* For pulsed tests, rest time of 20 min between pulses.

The function of the differentially expressed genes was predicted using GO analysis to obtain the most relevant biological process (BP), molecular function (MF) and cellular component (CC) categories activated in the full set of up- and down-regulated genes. Panther software was used to highlight the top 10 most enriched GO terms based on the gene signature from control *vs* free DOX and control *vs* mLNVs-DOX plus MH groups (Table S2 and S3). Comparative analysis revealed that GO categories in both conditions are similar. The enriched functions are mainly involved in the “cellular process” (BP), “binding” (MF) and “cellular anatomical entity” (CC). Moreover, the detailed Kyoto Encyclopedia of Genes and Genomes (KEGG) pathway enrichment results are shown in Tables S4 and S5 which highlight the top 10 enriched pathways in up- and down-differentially expressed genes. There was no significant differential activation of signaling pathways among the two groups. Among them, downregulated genes were mainly enriched for the “*wnt* signaling pathway” in both groups. The *wnt* signaling pathway regulates several cellular processes such as proliferation, survival, polarity, migration and cell differentiation.^54^ *Wnt* signaling is frequently abnormally activated in cancer cells.^55^ Furthermore, a meta-analysis based on the data from 1805 lung cancer patients reported a similar overexpression in *wnt* pathway genes with an inverse correlation to the overall survival of these patients.^56^

Upregulated genes were significantly enriched in the “*wnt* signaling pathway” for the control *vs* free DOX group while “gonadotropin-releasing hormone receptor pathway (*GnRH*)” and “inflammation mediated by chemokine and cytokine signaling pathway” were enriched for control *vs* mLNVs-DOX plus MH group. Inflammatory cytokines and chemokines are associated with cancer progression, and metastasis as well as angiogenesis,^57^ which makes them ideal antitumor agents or targets.^58^ On the other hand, the expression of *GnRH* has been shown to reduce cell invasion *in vitro* and inhibit metastasis *in vivo* across breast, ovarian and endometrial cancer cells.^59^ Additionally, DOX treatment has been found to induce apoptosis in GnRH receptor-positive human pancreatic cancer cells,^60^ while GnRH overexpression can trigger apoptosis through the activation of the Bcl-2/Bax/caspase pathway in human pancreatic cancer cells.^61^ *Bcl-2* and *Bax* are key apoptotic factors involved in cell apoptosis and autophagy processes.^62, 63^ Interestingly, in the comparison between the control untreated group and the mLNVs-DOX plus MH group, 11 up-DEGs belonging to the “apoptosis signaling pathway” were also enriched, namely, *Rax, Bclsl11, Hspa2, Fas, Hspa1l, Atf3, Ltb, Fos, Jun, Tnfrsd10b* and *Relb.* Overall, these findings suggest the potential involvement of the apoptosis signaling pathway in the observed synergistic effects of the combined treatment approach.

To further define the expression profiles of the melanoma cells after mLNVs-DOX plus MH treatment, our analysis was then focused on the statistically and significantly differentially expressed genes as shown in Figure 5. In this graph, genes that display statistically significant differential expression are highlighted in green and red, respectively. The most up- and down-regulated genes are highlighted in the plot. The genes *Osgin1, Mdm4, H1fO* and *Amotl2* were among the top significantly down-regulated. *Osgin1* is an apoptotic regulator gene, under the transcriptional control of p53,^64^ which could elevate cellular reactive oxygen species (ROS) production under stress conditions.^65^ In response to intra- and extra-cellular stress, the p53 pathway is activated to induce cell cycle arrest, cellular senescence and apoptosis to cease the propagation of mutations.^66^ Reduced *H1fO* expression is associated with uncontrolled self-renewal during tumor growth, whereas cells re-expressing *H1fO* acquire an epigenetic state that restricts their proliferative potential.^67^ *AMOTL2* is involved in low tumor proliferation, migration and invasion by regulating β-catenin nuclear localization, a key molecule of the *Wnt* signaling pathway.^68^ In turn, *Mdm4* is one of the major regulators of p53 expression and function, and can directly inhibit its transcriptional activity.^66^ *Mdm4* amplification and overexpression have been shown in many cancer types, such as melanoma, breast, glioma and soft tissue sarcoma.^66^ Nevertheless, our analysis revealed that *Mdm4* was one of the most significantly down-regulated genes in melanoma cells after mLNVs-DOX plus MH treatment compared to control cells. These data suggest that *Mdm4 may* contribute to the success of melanoma treatment by chemo-/thermotherapy using mLNVs. In line with these findings, four of the statistically significant up-regulated genes are related to tumor suppression: *Btg2*,^69^ *bcl2l11*,^70, 71^ *Cdkn1a*,^72–74^ *Fam46c*,^75, 76^ highlighting their involvement in the low survival of the cancer cells after mLNVs-DOX plus MH treatment. Other genes that were also strongly up-regulated (*Edar2*,^77^ *Polr2a*,^78–80^ *Cep170b*,^81, 82^ *Slc19a2*,^83, 84^ *Ulk1*^85^ and *Carhsp1*^86^) are related to cancer development and progression. These genes represent potential targets for follow-up innovative combined therapeutic approaches. However, it is important to consider the possibility of cross-talk between signaling pathways, which may limit the efficacy of these approaches. Inhibition of one pathway often leads to compensatory activation of another interconnected pathway, allowing for therapeutic escape.^87, 88^ Therefore, further experimental studies are required to elucidate the precise functions and clinical significance of all identified key genes in melanoma cancer in the future.

**Figure 5.**
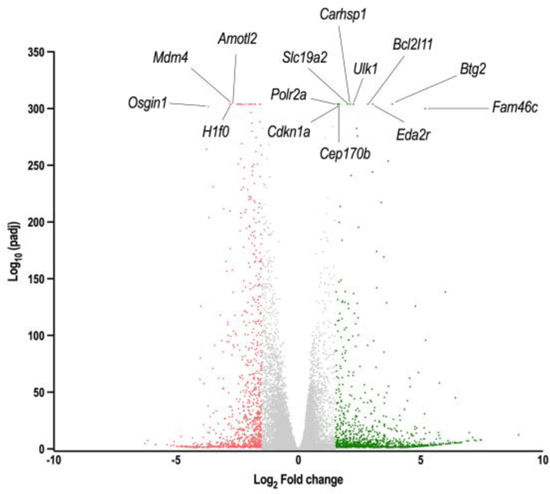
Volcano plot of differentially expressed genes based on RNA-seq analysis of untreated control melanoma cell *vs* mLNVs-DOX + MH therapy. Each gene is represented by a dot in the graph and the most differentially expressed up-and down-regulated genes are labeled in each plot. The x-axis and y-axis represent the log2 value of the fold change and the *t*-statistic as -log10 of the *p*-value, respectively. The genes represented in red (down-regulated) and green (up-regulated) are differentially expressed genes with > two-fold changes and a *p-*value ≤ 0.05, compared to the control.

As highlighted in the introduction, despite significant advancements in cancer treatment, the development of acquired resistance to chemotherapeutic drugs remains a major hurdle in patient care and overall treatment effectiveness.^27^ DOX, an intercalating agent that blocks DNA synthesis and transcription, ultimately leading to apoptosis,^89^ can cause genomic DNA alterations that contribute to the activation of biological processes associated with drug resistance.^90^ To explore the genes commonly altered in drug-resistant cells, a comparison was made between a list of drug resistance genes downloaded from NCBI^91^ and gene expression profiles of all tested groups compared to the control. Figure 6A highlights the most frequently altered genes (Table S6 provides a comprehensive description of these genes). This analysis identified a ten gene common signature (*Tlr4, Tlr2, Uvrag, Nlrp3, Bcl2, Prkaa1, Dpp4, Ppara, Fas, Vegfa*) correlated with drug resistance in cells treated with free DOX, while only six genes (*Ppara, Fas, Vegfa Nos3, Cdkn1a Icam1*) were associated with drug resistance in the mLNVs-DOX + MH group. The altered gene expression signature identified can in fact refect the biological response of cells in the presence of DOX (Figure S2), providing further support for our hypothesis that mLNVs-DOX combined with MH has the potential to prevent activation of drug resistance mechanisms.

**Figure 6.**
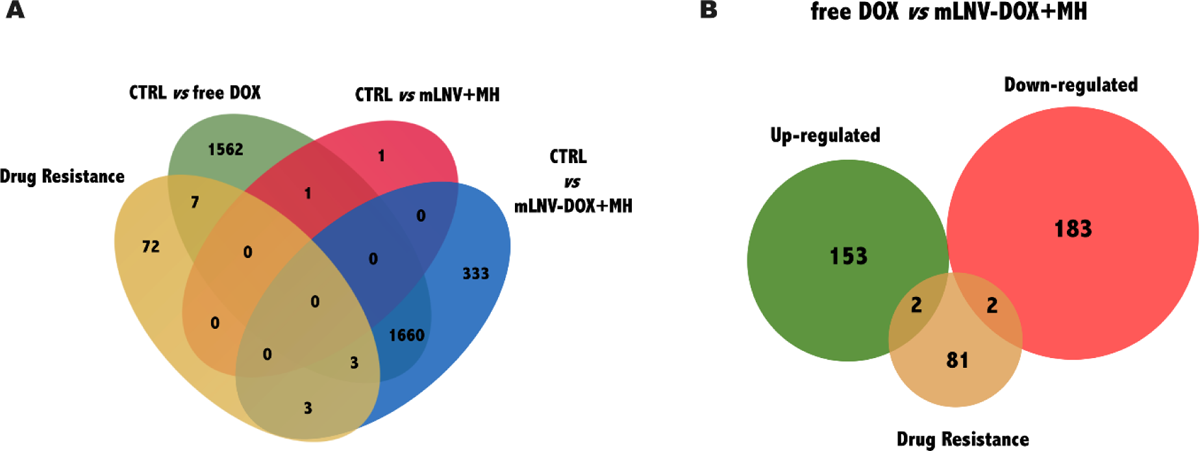
**(A)** Venn diagram of common signature among the drug resistance gene list (downloaded from NCBI) and the statistically significantly (FC ± 2 and *p-*value < 0.05) expressed genes of the tested groups compared to control. **(B)** Venn diagram was used to emphasize the common signature among the DR gene list and the statistically significantly up- and down-regulated genes on free DOX *vs* mLNVS-DOX+MH group. Circle size in B is only intended as a guide to the eye.

Moreover, the up- and down-regulated DEGs between free DOX *vs* mLNVs-DOX + MH and the drug resistance gene list were also overlapped and highlighted in Figure 6B. This overlapping analysis identified *Tlr2* and *Fas* (up-regulated), and *Nlrp3* and *Pdgfb* (down-regulated) as important genes involved in DOX drug resistance. Interestingly, the innate immune receptor Toll-like receptor 2 (*Tlr2*) is a key regulator of oncogene-induced senescence (OIS) and the senescence-associated secretory phenotype (SASP).^92^ Overexpression of *Tlr2* inducted a tumor suppressor response that impaired non-small cell lung cancer progression.^93^ In addition, *Fas* has been described as a key controller of the physiological regulation of programmed cell death (apoptosis).^94, 95^ The lack of expression of *Fas* is correlated with poor prognosis in melanoma tumors.^96^ Another gene highlighted in the drug resistance analysis is *Nlrp3* whose activation in melanoma cells drives cancer progression in mice.^97^ In addition, *Pdgfb* participates in different processes in solid tumors, including autocrine stimulation of cancer cell growth, recruitment of stroma fibroblasts and stimulation of local angiogenesis.^98^ Its downregulation significantly inhibits tumor progression and angiogenesis.^99^ Overall, the identified gene expression signature of free DOX *versus* mLNVs-DOX + MH with regards to drug resistance, further supports observed enhanced cell response after the combinatorial treatment in terms of viability data, highlighting the therapeutic potential of combining chemotherapy, mLNVs and hyperthermia treatment.

### In vivo assessment

Following the *in vitro* findings, the impact of pre-incubation time of mLNVs-DOX prior to AMF application was assessed in an *in vivo* setting. Adhering to the principles of the 3Rs policy, it was not feasible to test all the time points that were assessed in *in vitro*. Considering the pre-incubation, or circulation time in an *in vivo* scenario, two time points were chosen for evaluation: a short accumulation time of 3 h and a longer accumulation of 24 h. The 0 h time point, despite being optimal *in vitro*, was excluded as it was deemed necessary to allow some time for the mLNVs-DOX to reach and accumulate within the tumor environment. This decision ensured a more realistic representation of the treatment dynamics and allowed for a comprehensive assessment of the therapy’s effectiveness *in vivo*. Three doses were administered intravenously to each animal (1 every 48 h) followed every time by an AMF application 3 or 24 h after injection. Tumor growth was monitored daily for a total of 7 days (Fig 7A). Results both of tumor growth and final tumor weight at the end of the experiment (Fig 7B) show first, that the mLNVs-DOX are more efficient than free DOX and control mLNVs nanocomposites in controlling tumor growth. Also, the combination of mLNVs-DOX plus MH enhanced the effects of the therapy and helped reduce tumor growth further. Regarding the incubation/circulation time, both the 3 h and 24 h groups exhibited enhanced responses compared to the groups receiving free DOX and control mLNVs with MH. Animals that received AMF treatment 3 h after injection displayed the smallest tumor growth, indicating a more effective therapeutic outcome. However, the difference between the 3 h and 24 h groups, while significant, was not as substantial as anticipated based on the *in vitro* results. Interestingly, the *in vitro* tests showed that the combined chemo-/thermo-therapy had an equivalent effect to free DOX when applied after 24 h. However, in the *in vivo* setting, the results at 24 h exhibited a significant improvement (13.4%) compared to the free drug. These findings suggest that 24 h post-injection, most of the DOX was already released from the mLNVs, and the hyperthermia treatment no longer had a significant influence on the release process. Consequently, the therapeutic effect observed at the 24-hour time point *in vivo* was closer to the outcome of the mLNVs-DOX without hyperthermia group.

**Figure 7.**
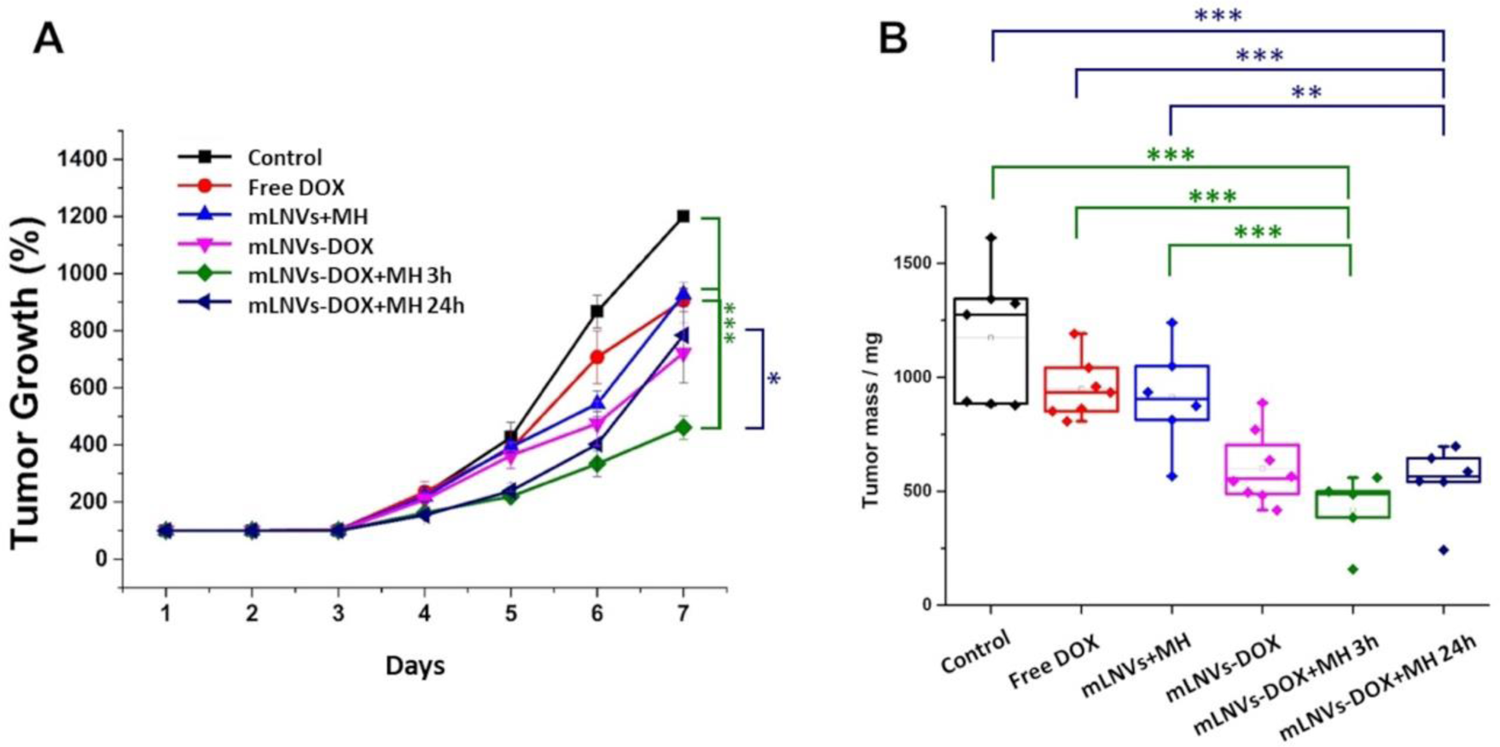
**(A**) Tumor growth of the different groups during the 7 days of the experiment. (**B**) Tumor weight of the different groups in the last day (7) of the tests. Box plot showing the average, percentile 25 and 75 (box), outliers (whiskers) and individual values (dots) showing significantly smaller tumor sizes of the mLNVs DOX with MH groups compared to the other groups (*** = *t =* 0.999; ** = *t =* 0.995).

A shorter incubation/circulation time led to improved outcomes, but the magnitude of the effect observed *in vitro* did not fully translate into the *in vivo* setup. Additional factors and complexities within the tumor microenvironment might influence the therapeutic response and necessitate further investigation to better understand the optimal timing for achieving maximum treatment efficacy.

## CONCLUSIONS

As the importance of combinatorial treatments increases in the fight against cancer, and so does their complexity and interrelationship, it is clear that the planning of the administration and follow-up processes requires more careful attention. The success of combinatorial treatments hinges on various factors, such as the compatibility of different therapies, optimal dosing, timing and sequence of administration. Each component contributes to the overall therapeutic outcome and patient response. Therefore, a comprehensive understanding of the interplay between different treatments and their effects becomes essential in order to maximize their potential benefits. In this work, we have explored the impact of the ‘where’, the ‘how’ and the ‘when’ of the application of magnetic hyperthermia as thermotherapy in combination with standard chemotherapy, on the outcome of the treatment. From the three parameters, the ‘when’ was shown to have the strongest effect on tumor growth/inhibition, and its importance could be traced back to physicochemical as well as biochemical parameters. The *in vitro* release profile of the chemotherapeutic drug marks the application protocol of the magnetic hyperthermia treatment. Relatively long incubation times (12-24 h) between NP administration and MH application, severely dampen the cytotoxic effects of the therapy *in vitro* to the point where no effects are observed. However, MH application shortly after NP administration synergistically enhances the therapeutic outcome. These same observations were reproduced *in vivo* in a preclinical mouse model, although differences were more subtle.

Moreover, looking into the transcriptomic profile of the combinatorial treatment and comparing it to the individual treatments, a couple of conclusions stand out. First, mild hyperthermia on its own does not seem to be suited for the treatment of melanoma as the aggressiveness of the cancer cells seems to increase. Second, differences in the therapeutic outcome with the combinatorial treatment might stem, at least in part, from the ability of the chemo/thermo-treatment to overcome resistance mechanisms, as stated by the changes in expression of a lower number of drug resistance genes in chemo/thermo-therapy when compared to chemotherapy alone. Besides drug resistance, the expression of other genes was highlighted in this analysis as potential responsible and therapeutic targets for an improved outcome. Overall, the transcriptomic analysis shed light on the complex interplay between different treatment modalities and gene expression patterns. It emphasizes the importance of understanding the molecular mechanisms underlying the therapeutic response and identifying novel targets to enhance treatment outcomes in cancer patients.

## METHODS

### mLNVs synthesis

A modified melt emulsification method was used for the preparation of mLNVs.^31^ In a standard preparation, 100 mg of Carnauba wax (Carnauba wax T1 pharmaceutical grade was a generous gift from Koster Keunen Holland BV - Raambrug 3, 5531 AG Bladel, The Netherlands) were mixed in a glass vial with a chloroform dispersion of Fe_3_O_4_ nanoparticles (prepared following a standard co-precipitation protocol)^100^ containing 30 mg of Fe. To this solution, 125 μL of a chloroform solution of DiO (3,3′-dioctadecyloxacarbocyanine perchlorate, 1 mg/mL) were added followed by a chloroform solution of doxorubicin (DOX, 20 mg, 1 mL). This mixture was heated under a heat gun until all the chloroform had evaporated and the wax melted. At this point, 2.25 mL of milliQ water followed by 0.25 mL of a water solution of Tween80 (50 mg/mL) were added to the vial and the sample was ultrasonicated for 2 min at 25% power at 20 s working intervals. Immediately after the sonication, the vial was immersed in ice to solidify the lipid nanoparticles. Once cold, the formulation was centrifuged (3000 rpm, 10 min), the pellet discarded, and the supernatant freeze dried in the presence of sucrose (0.9% w/w) as cryo-protectant.

### mLNVs characterization

The full characterization of the mLNVs formulation has been reported in detail in previous publications.^16, 101^

### Cell culture

B16F10 murine malignant melanoma cells (ATCC CRL-6475) were grown in Dulbecco’s Modified Eagle’s Medium (DMEM) supplemented with 10% fetal bovine serum and 1% antibiotic and were incubated at 37 °C under a 5% CO_2_ atmosphere.

### General magnetic hyperthermia application

4 × 10^5^ B16F10 cells were cultured in a 35 mm^2^ plate for 24 h and then exposed to 0.5 μg DOX/mL mLNVs formulations for different times (see Table 1). Then, the plate was placed in a live cell alternating magnetic field exposure system which has an ergonomic design and enables physiological temperature control and a 5% CO2 atmosphere (NanoTherics) and cells were exposed to the AMF for different times and under different application conditions (magnetic field intensity, frequency and mode) as shown in Table 1. At 48 h after the initial exposure to the mLNVs, the cells were washed twice with Hank’s balanced salt solution and standard cell viability studies were performed.

To test the spatial control of the magnetic hyperthermia procedure, a bespoke magnetic hyperthermia holder was designed, and 3D printed in PLA. The holder (Fig S1) had three different chambers placed at increasing distance from the coil center where glass slides were housed and used to grow the melanoma cells.

### Cell viability assays

Cell viability on the spatial control of magnetic hyperthermia was qualitatively evaluated using the cell-permeant dye Calcein AM, and images of live cells were acquired using a Nikon Eclipse Ti-E microscope. Melanoma cells were seeded in coverslips (25 mm) in a 6w plate at a density of 3.5× 10^5^ cells per well and left to grow overnight at 37 °C under 5 % CO_2_. The coverslips were then passed to the bespoke holder and the mLNVs-DOX (0.5 μg DOX/mL) were added to each compartment. Right after the addition of the nanoparticles, the holder was placed in the MH equipment where cells were treated for 1 h at 20 mT and 224 KHz. After treatment, the holder was closed and kept in the incubator to allow the cells to grow for 48 h at 37 °C/5 % CO_2_. Cells were then stained with 2 µM Calcein-AM for 35 min at room temperature and images of the coverslips were acquired one at a time.

Cell death was quantitatively evaluated using the Trypan blue exclusion assay after 48 h of NP exposure. Cultures were stained with Trypan Blue (10%), and live (bright) and dead (blue) cells were counted with a Neubauer’s chamber under a light microscope.

### TEM analysis

Cells were fixed with 3% glutaraldehyde in 0.12 M phosphate buffer and post-fixed in 1% buffered osmium tetroxide, dehydrated in a graded acetone series and embedded in Araldite. Then samples were sectioned (0.06 µm) with an ultramicrotome and stained with uranyl acetate. Images were acquired using a JEOL JEM 1011 microscope operated at 80 kV.

### Total RNA extraction and RNA-seq data processing

Total RNA was extracted from the cells using the RNeasy Mini Kit (Qiagen) according to the manufacturer’s instructions. Four groups were studied: Group 1: Control cells; Group 2: Cells treated with free DOX; Group 3: Cells treated with mLNVs + MH; Group 4: Cells treated with mLNVs-DOX + MH. To perform the RNA extraction, melanoma cells were collected 48h after initial exposure to the NPs. MH conditions: 224 kHz, 20 mT, 1 h at 37 °C and 5% CO2, DOX at 0.5 μg/mL.

Total RNA sequencing was performed by Novogene which also provided a preliminary analysis of the data. Benjamini and Hochberg’s test, with false discovery rate (FDR) was applied to identify statistically significant alterations in control *vs* treated samples. Alterations with Log2 FoldChange below 2 and *p* values above 0.05 were discarded. Raw and analyzed datasets have been deposited in NCBI’s Gene Expression Omnibus database and are accessible through GEO series accession number GSE234843.

### Functional Annotation

Gene function and GO and pathway enrichment analysis was annotated based on Uniprot^102^ and Pantherdb^103^ and Gene Ontology (GO) online resources.^104, 105^ The functional annotations were all determined based on the highest sequence similarity in these databases. GO enrichment analysis of differentially expressed genes was performed using standard GO Terms from the Gene Ontology Resource and a Fisher’s exact test with FDR *p*-value < 0.05 to estimate the statistical significance of the enrichment.

### Animal studies

Animals were maintained, handled, and sacrificed following the directive 2010/63/UE. C57BL/6 mice (8 – 10 weeks old) were housed with a 12 h light/dark cycle with free provision of food and water at the Experimentation Service (SEEA) of the University of Cantabria (CEA ES390750000849). Animal experimentation was approved by the local authority ‘Consejería de Medio Rural, Pesca y alimentación’ project number: PI-09-16.

### Tumor induction

A total of 5 × 10^5^ B16F10 melanoma cells in 100 μL DMEM were subcutaneously injected into the interscapular region of the mice. After 7 days tumors were palpable.

### Evaluation of anti-tumor and hyperthermia effects

When the tumor masses were palpable (ca. 7 days) the mice were randomly divided into 7 groups (*n* = 7 per group) as per Table 2.

**Table 2.** Experimental conditions for the *in vivo* tests.

| Group | Condition | Treatment regimen | Pre-incubation time before MH (h) |
| --- | --- | --- | --- |
| 1 | Control | Intravenous injection once every other day for three doses of saline solution | -- |
| 2 | Free DOX | Intravenous injection once every other day for three doses at a concentration of 2.5 mg DOX kg<sup>-1</sup> | -- |
| 3 | mLNVs-DOX | Intravenous injection once every other day for three doses at a concentration of 2.5 mg DOX kg<sup>-1</sup> | -- |
| 4 | mLNVs + MH | Intravenous injection once every other day for three doses at a conc equivalent to 2.5 mg DOX kg<sup>-1</sup> | 3 |
| 5 | mLNVs-DOX + MH | Intravenous injection once every other day for three doses at a concentration of 2.5 mg DOX kg<sup>-1</sup> | 3 |
| 6 | mLNVs + MH | Intravenous injection once every other day for three doses at a conc equivalent to 2.5 mg DOX kg<sup>-1</sup> | 24 |
| 7 | mLNVs-DOX + MH | Intravenous injection once every other day for three doses at a concentration of 2.5 mg DOX kg<sup>-1</sup> | 24 |

After the administration (3 h for groups 4 and 5, 24 h for groups 6 and 7), mice in groups 4, 5, 6 and 7 were anesthetized by inhalation (1% isoflurane), placed in a water-cooled induction coil (50 mm internal diameter, nine turns) with the tumors located in the center of the coil and exposed to 174.5 kHz, 23 mT for 1 h. The body weight and tumor size of each animal were monitored daily. The estimated tumor volume was calculated using the following formula:

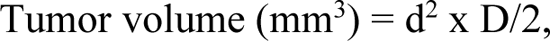

where d and D are the shortest and longest diameter in mm, respectively. At the end of the experimental period, the mice were euthanized, and their tissues were dissected and harvested.

### Statistical analysis

Groups were compared using ANOVA tests. In cases where ANOVA yielded significant results, pairwise comparisons and Student’s *t*-tests were performed. The statistical analyses were conducted using SPSS, version 19.0.

## ACKNOWLEDGEMENTS

The authors would like to acknowledge the financial support from the Portuguese Foundation for Science and Technology (FCT, Fundação para a Ciência e a Tecnologia) through projects NORTE-01-0145-FEDER-028052 (SELF-i), NORTE-01-0145-FEDER-031142 (MagTargetON) and PTDC/QUI-OUT/3143/2021 (UnTAM). We would also acknowledge the financial support from the Spanish Ministry of Science and Innovation through the project TED2021-129248 B–I00 funded by MCIN/AEI/10.13039/501100011033 and “European Union Next Generation EU/PRTR”; and the Spanish Instituto de Salud Carlos iii under Project ref. PI22/00030, the two projects co-funded by the European Regional Development Funds. LGH thanks the Agencia Estatal de Investigación for the Juan de la Cierva Incorporación Grant (IJC2020-043746-I).

## Supporting Information

Available composition of the mLNVs subject of the study, tables with the gene ontology results, DOX drug resistance expression signature, MH system used in the study, Cell viability of B16F10 cells after treatment, DOX release profile from mLNVs, Cell death of B16F10 cells after different pre-incubation times and under different operational parameters, Cell viability of B16F10 cells treated with mLNVs-DOX at two different concentrations, TEM images of B16F10 cells incubated with mLNVs-DOX, Distribution of gene expression levels among the different samples/conditions, Venn diagram of the top 20 altered genes,

