## Supporting information for "The critical role of spatio-temporal control in combinatorial chemo- and magnetic hyperthermia thermo-therapy: ‘the where’, ‘the how’ and ‘the when’"

[2]Grupo de Nanomedicina. Departamento de Biología Molecular, Universidad de Cantabria- Instituto de Investigación Valdecilla-IDIVAL, Herrera Oria s/n, 39011, Santander, Spain

[3]Departamento de Anatomía y Biología Celular, Universidad de Cantabria, Herrera Oria s/n, 39011, Santander, Spain

### Supporting information

**Table S1.** Composition of the mLNVs subject of this study.

| Component | Mass/Proportion |
|---|---|
| **Wax** | 100 mg |
| **Tween 80** | 12.5% |
| **Magnetic nanoparticles** | 30% |
| **DOX** | 20% |
| **DiO** | 0.125% |

**Table S2.** Gene Ontology enrichment of up- and down-regulated differentially expressed genes in CTRL *vs* free DOX.

| | **Biological Process** | | **Genes** | | **Biological Process** | | **Genes** |
|---|---|---|---|---|---|---|---|
| Up-regulated | cellular process | GO:0009987 | 465 | Down-regulated | cellular process | GO:0009987 | 749 |
| | biological regulation | GO:0065007 | 316 | | metabolic process | GO:0008152 | 501 |
| | metabolic process | GO:0008152 | 241 | | biological regulation | GO:0065007 | 483 |
| | response to stimulus | GO:0050896 | 174 | | response to stimulus | GO:0050896 | 178 |
| | signaling | GO:0023052 | 136 | | signaling | GO:0023052 | 149 |
| | multicellular organismal process | GO:0032501 | 117 | | localization | GO:0051179 | 122 |
| | localization | GO:0051179 | 111 | | developmental process | GO:0032502 | 86 |
| | developmental process | GO:0032502 | 101 | | multicellular organismal process | GO:0032501 | 85 |
| | immune system process | GO:0002376 | 37 | | immune system process | GO:0002376 | 25 |
| | biological adhesion | GO:0022610 | 31 | | biological adhesion | GO:0022610 | 19 |
| | **Molecular Function** | | **Genes** | | **Molecular Function** | | **Genes** |
| | binding | GO:0005488 | 293 | | binding | GO:0005488 | 458 |
| | catalytic activity | GO:0003824 | 178 | | catalytic activity | GO:0003824 | 244 |
| | molecular transducer activity | GO:0060089 | 77 | | transcription regulator activity | GO:0140110 | 168 |
| | transcription regulator activity | GO:0140110 | 75 | | molecular function regulator | GO:0098772 | 58 |
| | transporter activity | GO:0005215 | 67 | | molecular transducer activity | GO:0060089 | 34 |
| | molecular function regulator | GO:0098772 | 43 | | transporter activity | GO:0005215 | 26 |
| | ATP-dependent activity | GO:0140657 | 14 | | ATP-dependent activity | GO:0140657 | 16 |
| | cytoskeletal motor activity | GO:0003774 | 8 | | molecular adaptor activity | GO:0060090 | 8 |
| | structural molecule activity | GO:0005198 | 6 | | cytoskeletal motor activity | GO:0003774 | 7 |
| | molecular adaptor activity | GO:0060090 | 4 | | structural molecule activity | GO:0005198 | 7 |
| | **Cellular Component** | | **Genes** | | **Cellular Component** | | **Genes** |
| | cellular anatomical entity | GO:0110165 | 507 | | cellular anatomical entity | GO:0110165 | 705 |
| | protein-containing complex | GO:0032991 | 96 | | protein-containing complex | GO:0032991 | 202 |

**Table S3**. Gene Ontology enrichment of up- and down-regulated differentially expressed genes in CTRL *vs* mLNV-DOX+MH.

| | **Biological Process** | | **Genes** | | **Biological Process** | | **Genes** |
|---|---|---|---|---|---|---|---|
| **Up-regulated** | cellular process | GO:0009987 | 369 | **Down-regulated** | cellular process | GO:0009987 | 325 |
| | biological regulation | GO:0065007 | 256 | | biological regulation | GO:0065007 | 217 |
| | metabolic process | GO:0008152 | 196 | | metabolic process | GO:0008152 | 211 |
| | response to stimulus | GO:0050896 | 142 | | response to stimulus | GO:0050896 | 79 |
| | signaling | GO:0023052 | 109 | | signaling | GO:0023052 | 71 |
| | multicellular organismal process | GO:0032501 | 85 | | localization | GO:0051179 | 62 |
| | developmental process | GO:0032502 | 75 | | developmental process | GO:0032502 | 47 |
| | localization | GO:0051179 | 73 | | multicellular organismal process | GO:0032501 | 46 |
| | immune system process | GO:0002376 | 26 | | immune system process | GO:0002376 | 13 |
| | locomotion | GO:0040011 | 24 | | biological adhesion | GO:0022610 | 11 |
| | **Molecular Function** | | **Genes** | | **Molecular Function** | | **Genes** |
| | binding | GO:0005488 | 234 | | binding | GO:0005488 | 210 |
| | catalytic activity | GO:0003824 | 141 | | catalytic activity | GO:0003824 | 109 |
| | transcription regulator activity | GO:0140110 | 69 | | transcription regulator activity | GO:0140110 | 86 |
| | molecular transducer activity | GO:0060089 | 63 | | molecular function regulator | GO:0098772 | 21 |
| | transporter activity | GO:0005215 | 40 | | molecular transducer activity | GO:0060089 | 19 |
| | molecular function regulator | GO:0098772 | 32 | | transporter activity | GO:0005215 | 17 |
| | ATP-dependent activity | GO:0140657 | 13 | | ATP-dependent activity | GO:0140657 | 6 |
| | cytoskeletal motor activity | GO:0003774 | 7 | | molecular adaptor activity | GO:0060090 | 6 |
| | structural molecule activity | GO:0005198 | 4 | | cytoskeletal motor activity | GO:0003774 | 3 |
| | molecular adaptor activity | GO:0060090 | 2 | | translation regulator activity | GO:0045182 | 2 |
| | **Cellular Component** | | **Genes** | | **Cellular Component** | | **Genes** |
| | cellular anatomical entity | GO:0110165 | 375 | | cellular anatomical entity | GO:0110165 | 311 |
| | protein-containing complex | GO:0032991 | 67 | | protein-containing complex | GO:0032991 | 73 |

**Table S4** Pathways enrichment of up- and down-regulated differentially expressed genes in CTRL *vs* free DOX.

| | Pathways | | Genes |
|---|---|---|---|
| Up-regulated | Wnt signaling pathway | P00057 | 18 |
| | Gonadotropin-releasing hormone receptor pathway | P06664 | 17 |
| | Heterotrimeric G-protein signaling pathway-Gi alpha and Gs alpha mediated pathway | P00026 | 16 |
| | Inflammation mediated by chemokine and cytokine signaling pathway | P00031 | 15 |
| | Angiogenesis | P00005 | 14 |
| | Nicotinic acetylcholine receptor signaling pathway | P00044 | 14 |
| | Blood coagulation | P00011 | 12 |
| | TGF-beta signaling pathway | P00052 | 10 |
| | Integrin signalling pathway | P00034 | 10 |
| | Heterotrimeric G-protein signaling pathway-Gq alpha and Go alpha mediated pathway | P00027 | 10 |

| | Pathways | | Genes |
|---|---|---|---|
| Down-regulated | Wnt signaling pathway | P00057 | 26 |
| | Gonadotropin-releasing hormone receptor pathway | P06664 | 19 |
| | Angiogenesis | P00005 | 16 |
| | Inflammation mediated by chemokine and cytokine signaling pathway | P00031 | 16 |
| | CCKR signaling map | P06959 | 16 |
| | Apoptosis signaling pathway | P00006 | 13 |
| | Integrin signalling pathway | P00034 | 13 |
| | PDGF signaling pathway | P00047 | 13 |
| | Cadherin signaling pathway | P00012 | 13 |
| | p53 pathway | P00059 | 10 |

**Table S5** Pathways enrichment of up- and down-regulated differentially expressed genes in CTRL *vs* mLNV-DOX+MH.

| | Pathways | | Genes | | Pathways | | Genes |
|---|---|---|---|---|---|---|---|
| Up-regulated | Gonadotropin-releasing hormone receptor pathway | P06664 | 16 | Down-regulated | Wnt signaling pathway | P00057 | 13 |
| | Inflammation mediated by chemokine and cytokine signaling pathway | P00031 | 16 | | Gonadotropin-releasing hormone receptor pathway | P06664 | 12 |
| | Angiogenesis | P00005 | 12 | | Angiogenesis | P00005 | 8 |
| | Wnt signaling pathway | P00057 | 12 | | Inflammation mediated by chemokine and cytokine signaling pathway | P00031 | 7 |
| | Apoptosis signaling pathway | P00006 | 11 | | Heterotrimeric G-protein signaling pathway-Gq alpha and Go alpha mediated pathway | P00027 | 7 |
| | Alzheimer disease-presenilin pathway | P00004 | 11 | | CCKR signaling map | P06959 | 7 |
| | Heterotrimeric G-protein signaling pathway-Gi alpha and Gs alpha mediated pathway | P00026 | 11 | | Apoptosis signaling pathway | P00006 | 6 |
| | Integrin signalling pathway | P00034 | 10 | | TGF-beta signaling pathway | P00052 | 6 |
| | Huntington disease | P00029 | 9 | | PDGF signaling pathway | P00047 | 6 |
| | TGF-beta signaling pathway | P00052 | 7 | | Cadherin signaling pathway | P00012 | 6 |

**Table S6**. Doxorubicin drug resistance expression signature.

| | | Gene | Gene Description | FC | *padj* | Main Function | Potential effect* |
|---|---|---|---|---|---|---|---|
| CTRL *vs* free DOX | | *Tlr4* | Toll-like receptor 4 | -1,72 | 1,34E-80 | The receptor encoded by this gene mediates the innate immune response to bacterial lipopolysaccharide through synthesis of pro-inflammatory cytokines and chemokines. | - |
| | | *Tlr2* | Toll-like receptor 2 | -2,37 | 1,46E-03 | Plays a fundamental role in pathogen recognition and activation of innate immunity. | - |
| | | *Uvrag* | UV radiation resistance associated gene | -1,79 | 0,00E+00 | *Uvrag* overexpression enhances autophagy and reduces proliferation, suggesting that controls cell growth by regulating autophagy. | - |
| | | *Nlrp3* | NLR family, pyrin domain containing 3 | 3,66 | 1,12E-02 | Promotes cancer growth and metastasis by instructing an immunosuppressive TME, mainly through IL-1β. | - |
| | | *Bcl2* | B cell leukemia/lymphoma 2 | 3,14 | 0,00E+00 | Abnormal expression and chromosomal translocations are associated with cancer progression. | - |
| | | *Prkaa1* | protein kinase, AMP-activated, alpha 1 catalytic subunit | -1,73 | 0,00E+00 | Involved in several processes, including regulation of gene expression; regulation of organelle organization; and regulation of protein modification process. | + |
| | | *Dpp4* | dipeptidylpeptidase 4 | -4,32 | 2,42E-02 | Participates in various physiological and pathological processes by regulating energy metabolism, inflammation, and immune function. DPP4 gene has been shown to be a tumor suppressor in other cancers, as melanoma and ovarian cancer. | - |
| | Common expressed genes | *Ppara* | Peroxisome proliferator-activated receptor alpha | -2,47 | 5,13E-13 | Is involved in several functions, including DNA-binding transcription factor activity; RNA polymerase II cis-regulatory region sequence-specific DNA binding activity; and lipid binding activity. | + |
| | | *Fas* | Tumor necrosis factor receptor superfamily member 6 | 3,01 | 2,20E-04 | Extrinsic apoptotic signalling pathway. | + |
| | | *Vegfa* | Vascular endothelial growth factor A | 1,75 | 7,93E-121 | Its expression is correlated with tumor stage and progression. | - |
| CTRL *vs* mLNVs-DOX+MH | | *Ppara* | Peroxisome proliferator-activated receptor alpha | -2,44 | 1,59E-12 | Is involved in several functions, including DNA-binding transcription factor activity; RNA polymerase II cis-regulatory region sequence-specific DNA binding activity; and lipid binding activity. | + |
| | | *Fas* | Tumor necrosis factor receptor superfamily member 6 | 4,70 | 1,95E-12 | Extrinsic apoptotic signalling pathway. | + |
| | | *Vegfa* | Vascular endothelial growth factor A | 2,42 | 1,24E-276 | Its expression is correlated with tumor stage and progression. | - |
| | | *Nos3* | Nitric oxide synthase 3 | -1,50 | 2,41E-04 | Inhibits apoptosis and promote angiogenesis, proliferation, invasiveness, and immunosuppression of malignant tumors. | + |
| | | *Cdkn1a* | Cyclin-dependent kinase inhibitor 1A (P21) | 1,63 | 0,00E+00 | Functions as a regulator of cell cycle progression at the G1 phase. Mice that lack this gene have the ability to regenerate damaged or missing tissue. | - |
| | | *Icam1* | Intercellular adhesion molecule 1 | -2,04 | 4,15E-02 | It participates in the innate immune response. | - |

*Oversimplified potential effect on the therapy. - potential effect, negative effect on the overall outcome of the therapy; +, positive effect on the overall outcome of the therapy.

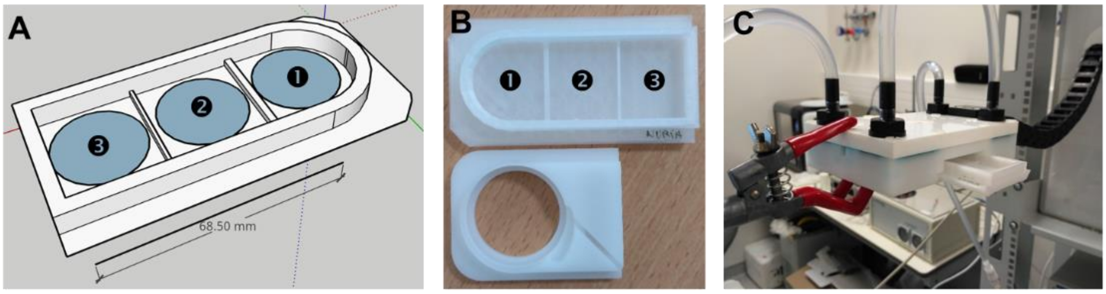


**Figure S1.** **A**, sketch of the bespoke holder designed for the local AMF tests. **B**, comparison of the standard commercial MH holder (bottom) and the bespoke one (top). **C**, *in vitro* AMF applicator enabling a controlled T and atmospheric conditions.

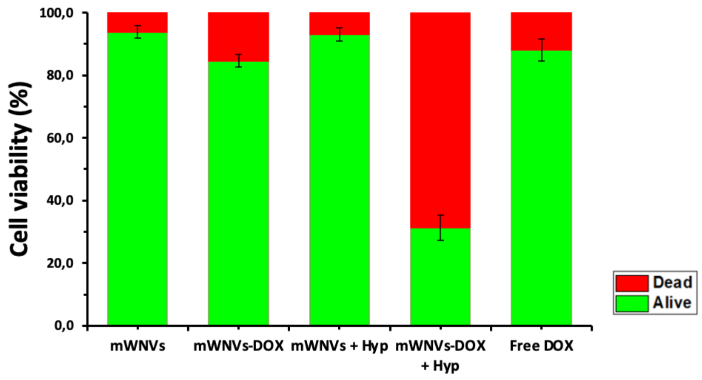


**Figure S2.** Cell viability of B16F10 cells after treatment (from left to right) with mLNVs, mLNVs-DOX, mLNVs+MH, mLNVs-DOX+MH and free DOX at a concentration of 0.5 μg DOX/mL or equivalent in those formulations without DOX.

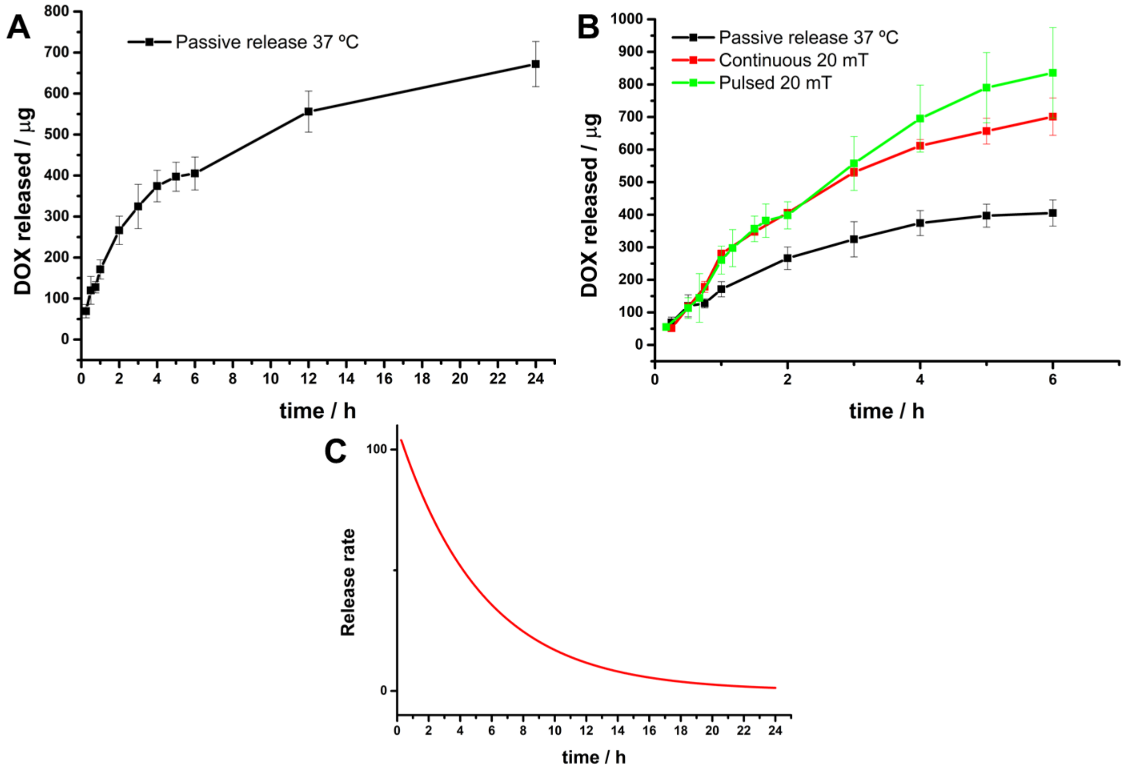


**Figure S3.** (**A**) DOX release profile under passive conditions at 37 ºC along 24 h. (**B**) DOX release profile under passive conditions at 37 ºC (black), continuous MH at 20 mT and 224 kHz (red) and pulsed MH at 20 mT and 224 kHz (green) during the first 6 h of release. MH (continuous or pulsed) was applied during the first 3h. Pulsed protocol: 20 mT, 15 min ON, 15 min off. (**C**) DOX release rate calculated through the first derivative of the exponential growth fitting of the passive release in A.

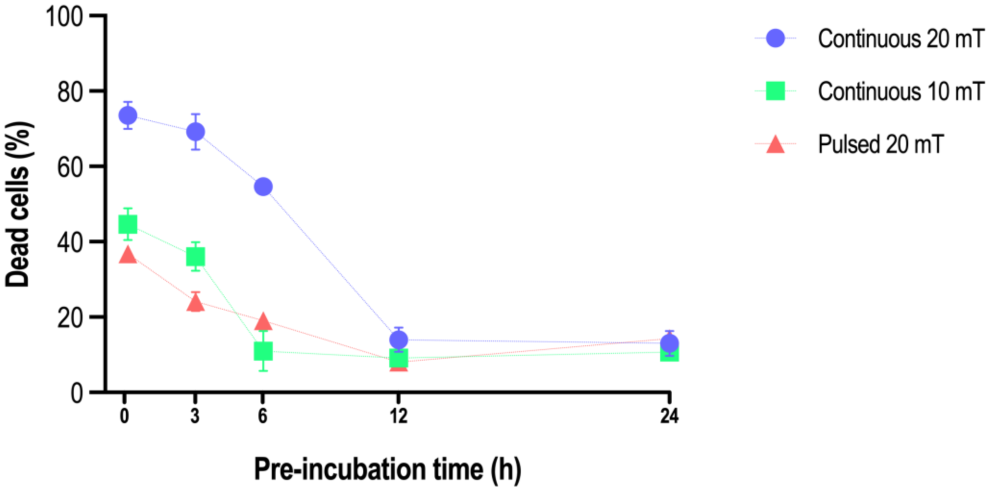


**Figure S4.** Cell death of B16F10 cells after pre-incubation with mLNVs-DOX (0.5 μg DOX/mL) for different times (0 to 24 h) +MH (continuous for 1h at 224 kHz and 20 mT; continuous for 1h at 224 kHz and 10 mT; pulsed 3x 10 min + 20 min rest, at 224 kHz and 20 mT).

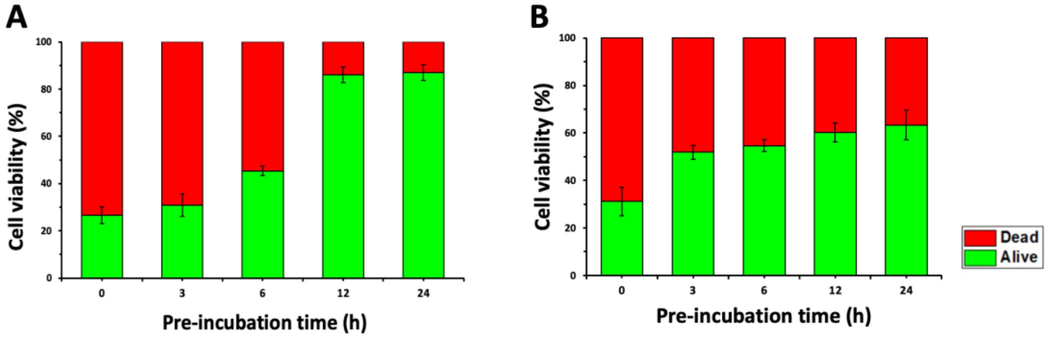


**Figure S5.** Cell viability of B16F10 cells treated with mLNVs-DOX at **(A)** 0.5 μg DOX/mL and (**B)** 1.0 μg DOX/mL and application of AMF at 20 mT for 1h after 0, 3, 6, 12 and 24 h of pre-incubation time.

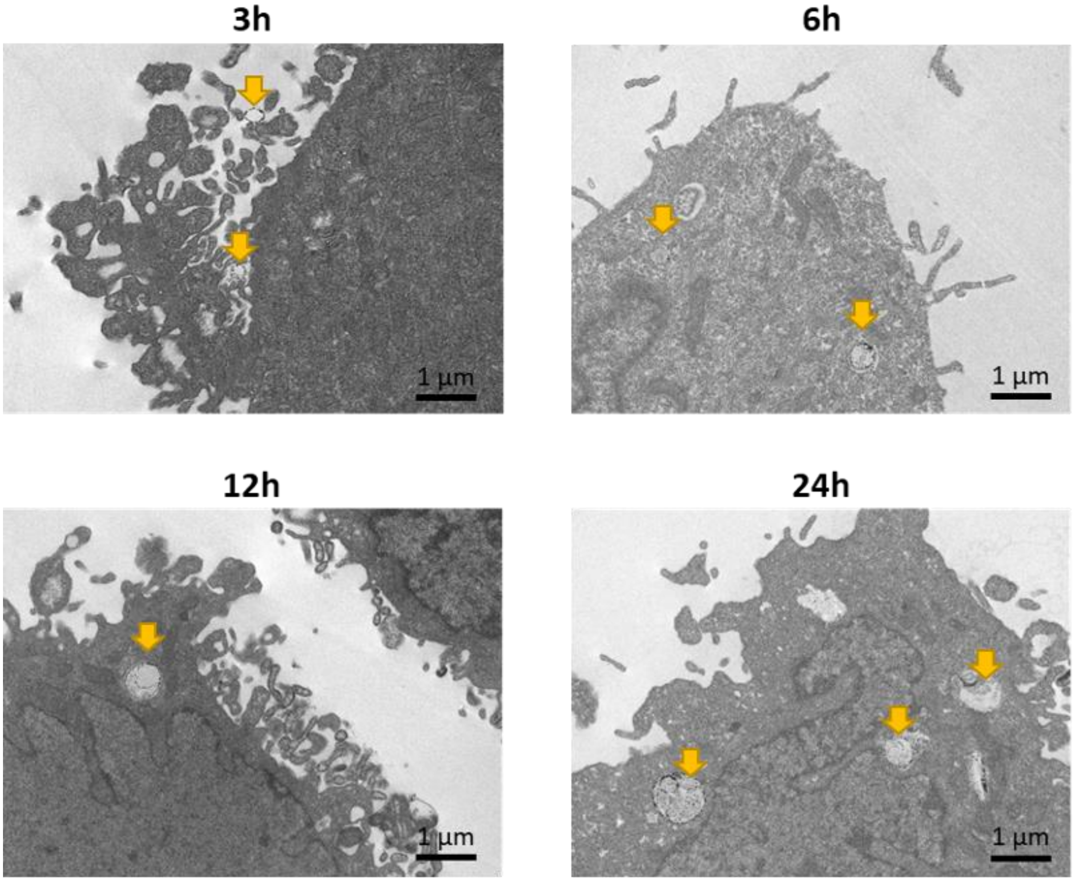


**Figure S6.** TEM images of B16F10 cells incubated with mLNVs-DOX (0.5 µg DOX/mL) for different times. Visible mLNVs (yellow arrows) are interacting with the membrane (3h) or inside the cell (6, 12 and 24h).

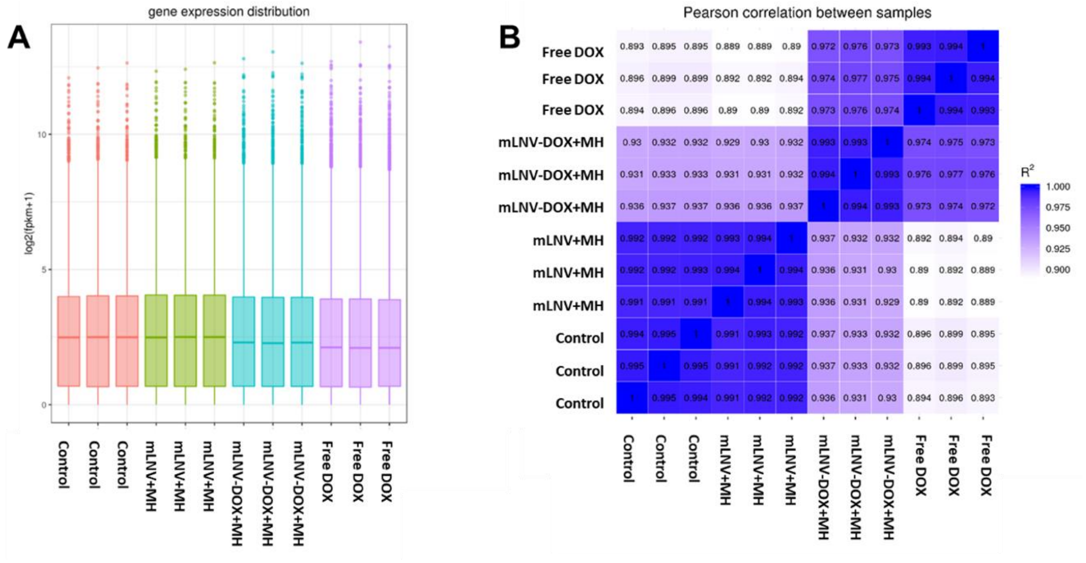


**Figure S7. (A)** Distribution of gene expression levels among the different samples/conditions. Parameters of the box plots include maximum, upper quartile, mid-value, lower quartile and minimum. The plot shows that gene expression levels do not change significantly between conditions. **(B)** Correlation coefficient matrix ($R^2$ = square of Pearson correlation coefficient (R)), confirming the reliability of the data.

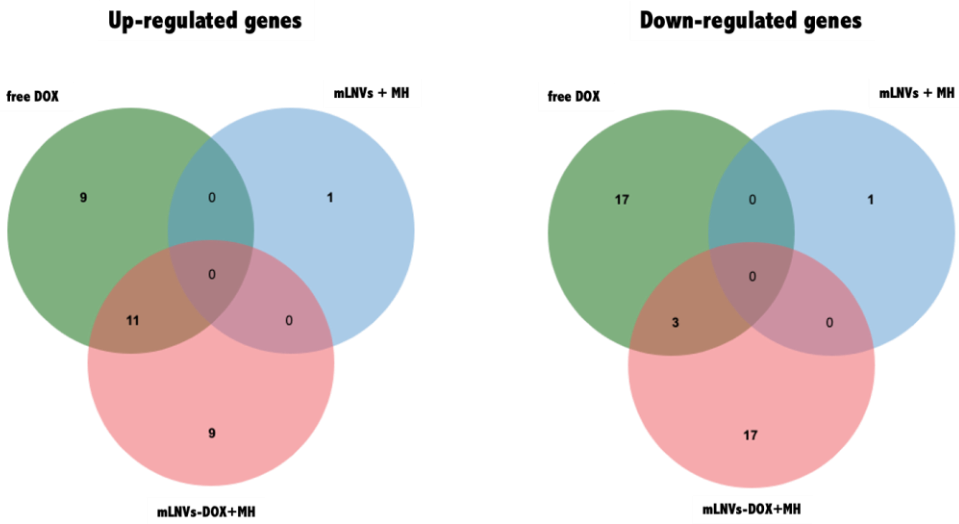


**Figure S8.** Venn diagram of the top 20 altered abundant genes as effect of chemo- and/or thermo-therapy *versus* control cell, considering the cut-off FC $\pm 2$ and p-value $< 0.05$.
